# Membrane contact site detection (MCS-DETECT) reveals dual control of rough mitochondria-ER contacts

**DOI:** 10.1101/2022.06.23.497346

**Authors:** Ben Cardoen, Kurt Vandevoorde, Guang Gao, Parsa Alan, William Liu, Ellie Tiliakou, A. Wayne Vogl, Ghassan Hamarneh, Ivan R. Nabi

**Affiliations:** School of Computing Science, Simon Fraser University, Burnaby, BC, Canada V5A 1S6; Department of Cellular & Physiological Sciences, Life Sciences Institute, University of British Columbia, Vancouver, BC, Canada V6T 1Z3; School of Biomedical Engineering, University of British Columbia, Vancouver, BC, Canada V6T 1Z3

## Abstract

Identification and morphological analysis of mitochondria-ER contacts (MERCs) by fluorescent microscopy is limited by sub-pixel resolution inter-organelle distances. Application of a Membrane Contact Site (MCS) detection algorithm, MCS-DETECT, to 3D STED super-resolution image volumes reconstructs sub-resolution MERCs. MCS-DETECT shows that elongated ribosome-studded riboMERCs, present in HT-1080 but not COS-7 cells, are morphologically distinct from smaller smooth contacts and larger contacts induced by mitochondria-ER linker expression in COS-7 cells. riboMERC expression is reduced in Gp78 knockout HT-1080 cells and induced by Gp78 ubiquitin ligase activity in COS-7 cells. Knockdown of the riboMERC tether RRBP1 eliminates riboMERCs in both wild-type and Gp78 knockout HT-1080 cells. By MCS-DETECT, Gp78-dependent riboMERCs present complex tubular shapes that intercalate between and contact multiple mitochondria, that are lost upon RRBP1 knockdown. MCS-DETECT of 3D whole cell super-resolution image volumes therefore identifies a novel dual regulatory mechanism for tubular riboMERCs, whose formation is dependent on RRBP1 and size modulated by Gp78 E3 ubiquitin ligase activity.

**eTOC Summary:** Application of the sub-pixel resolution Membrane Contact Site (MCS) detection algorithm, MCS-DETECT, to 3D STED super-resolution image volumes identifies a novel dual regulatory mechanism for tubular riboMERCs, whose formation is dependent on RRBP1 and size modulated by Gp78 E3 ubiquitin ligase activity.

## Introduction

In the cell, organelles communicate with each other at membrane contact sites (MCS) where two membranes come in close proximity, as close as 10-30 nm, without fusing (Helle et al., 2013; Valm et al., 2017). Mitochondria-endoplasmic reticulum contacts (MERCs), an MCS subclass, are hubs for exchange between the endoplasmic reticulum (ER) and mitochondria, enabling calcium transfer required for mitochondrial enzyme activity and ATP production, phospholipid and sterol biosynthesis, mitochondrial dynamics and metabolism as well as the execution of cell death programs (Rowland and Voeltz, 2012). MERCs are closely associated with disease progression, including cancer, neurodegenerative, cardiovascular and other diseases (Barazzuol et al., 2021; Díaz et al., 2021; Markovinovic et al., 2022). MERCs were traditionally thought to involve smooth ER, i.e. membrane regions of the ER devoid of ribosomes, and represent close contacts (∼10-15 nm) between the two organelles (Goetz and Nabi, 2006); ribosome-studded rough ER riboMERCs (25 nm contact distance) are found in liver and called WrappER-associated mitochondria (WAM) as they wrap around mitochondria (Anastasia et al., 2021; Csordas et al., 2006; Giacomello and Pellegrini, 2016; Ilacqua et al., 2022). Wider (50-60 nm) riboMERCs were identified in metastatic HT-1080 fibrosarcoma cells and in HEK293 cells where they are regulated by the Gp78 E3 ubiquitin ligase and interaction between mitochondrial Outer Membrane Protein 25 (OMP25), also called Synaptojanin-2-binding protein (SYNJ2BP), and its ER partner, ribosome-binding protein 1 (RRBP1), respectively (Hung et al., 2017; Wang et al., 2015). Other studies have reported MERCs of varying distances ranging from 10-80 nm (Giacomello and Pellegrini, 2016) highlighting the diversity of MERCs.

The specificity of labeling and ability to study dynamic fluorescent tagged proteins in living cells makes fluorescent microscopy the method of choice to characterize the diversity, dynamics and molecular mechanisms underlying MERC formation. However, analysis of MERCs by fluorescence microscopy faces three major hurdles: 1) the distance between the ER and mitochondria is below the resolution of optical microscopy (200-250 nm) due to diffraction limits; 2) MERC segmentation approaches are sensitive to subjective parameter settings such that accurate thresholding of the ER, ranging from isolated peripheral tubules to the dense central ER matrix, is particularly challenging; 3) bifluorescent complementation systems of varying linker lengths present differential detection of MERCs, but may promote or stabilize MERC formation (Cieri et al., 2018; Harmon et al., 2017; Vallese et al., 2020). Earlier work using diffraction limited confocal microscopy showing that ER tubules mark mitochondrial constrictions (Friedman et al., 2011) has been confirmed using super-resolution single molecule localization microscopy (SMLM) (Shim et al., 2012) and live cell 2D stimulated emission depletion (STED) imaging (Bottanelli et al., 2016). 2D STED characterized roles for ER shaping proteins in control of peripheral ER tubule nanodomains and fenestrations in ER sheets (Gao et al., 2019; Schroeder et al., 2019). 3D super-resolution whole cell analysis by structured illumination (SIM) or 3D STED microscopy achieves ∼120 nm lateral and ∼250 nm axial resolution and identified tubular matrices in peripheral ER sheets and in the dense central ER of Zika virus infected cells (Long et al., 2020; Nixon-Abell et al., 2016). SIM and STED therefore represent optimal super-resolution imaging approaches to study the distribution and morphology of MERCs in 3D whole cell views.

However, the intervening space between ER and mitochondria remains far smaller than the resolution provided by SIM or 3D STED. In addition, detection of MERCs requires analytical approaches that can accurately assess overlap between two fluorescent channels acquired independently, facing challenges of varying background density and difficulty of thresholding signals of varying signal to noise ratio. Recently, the optimal transport distance between two fluorescence distributions was formulated as an alternative approach to colocalization (Tameling et al., 2021; Wang and Yuan, 2021). However, whether this method is able to recover interactions at sub-precision distances of MERCs is not known. By reducing the anisotropy using multiple unregistered samples and mathematical optimization, more precise colocalization is becoming possible (Fortun et al., 2018); however, this method does not directly quantify sub-precision interfaces. Quantitative detection of MERCs in 3D cell volumes for fluorescent microscopy therefore requires novel analytical approaches that apply sub-pixel resolution detection of interaction zones to whole cell super-resolution imaging approaches.

To this end we developed a multichannel differential analysis to reconstruct the interface at sub-pixel precision: MCS-DETECT. Without segmentation, the algorithm adapts robustly to the intensity variations between fluorescent channels and samples, resulting in highly sensitive detection independent of variations in local signal or background intensity differentials. Application to 3D STED super-resolution microscopy images of cells distinguishes different classes of MERCs and defines a novel dual mechanism that controls the formation of a distinct class of extended, convoluted, tubular riboMERCs. Formation of tubular riboMERCS is dependent on expression of the riboMERC tether RRBP1 and their size and shape modulated by Gp78 E3 ubiquitin ligase activity.

## RESULTS

### Distinct MERCs in COS-7 and HT-1080 cells

EM analysis shows the presence of elongated rough ER-mitochondria (RER-mito) contacts in HT-1080 cells, as previously reported (Wang et al., 2015) while COS-7 cells present predominantly shorter, smooth ER-mitochondria (SER-mito) contacts (Fig. 1). Further, SER-mito and RER-mito contacts co-exist in HT-1080 cells as a single unit, with a smaller smooth MERC extending from a more elongated RER-mito contact (Fig. 1A), as previously reported (Giacomello and Pellegrini, 2016; Wang et al., 2015). Defining RER-mito contacts as any contact site containing at least one ribosome within the interorganellar space, we report here a 55 nm spacing between ER and mitochondria of HT-1080 RER-mito contacts and 15 nm spacing of SER-mito contacts, corresponding to our previous study (Wang et al., 2015). In COS-7 cells, RER-mito contacts are ∼35 nm in width while SER-mito contacts are ∼25 nm in width (Fig. 1B). The different spacing of both classes of MERCs in HT-1080 and COS-7 cells highlights the varied spacing of MERCs in different cells and tissues (Csordas et al., 2006; Giacomello and Pellegrini, 2016; Hung et al., 2017; Parlakgül et al., 2020; Wang et al., 2015).

**Figure 1.**
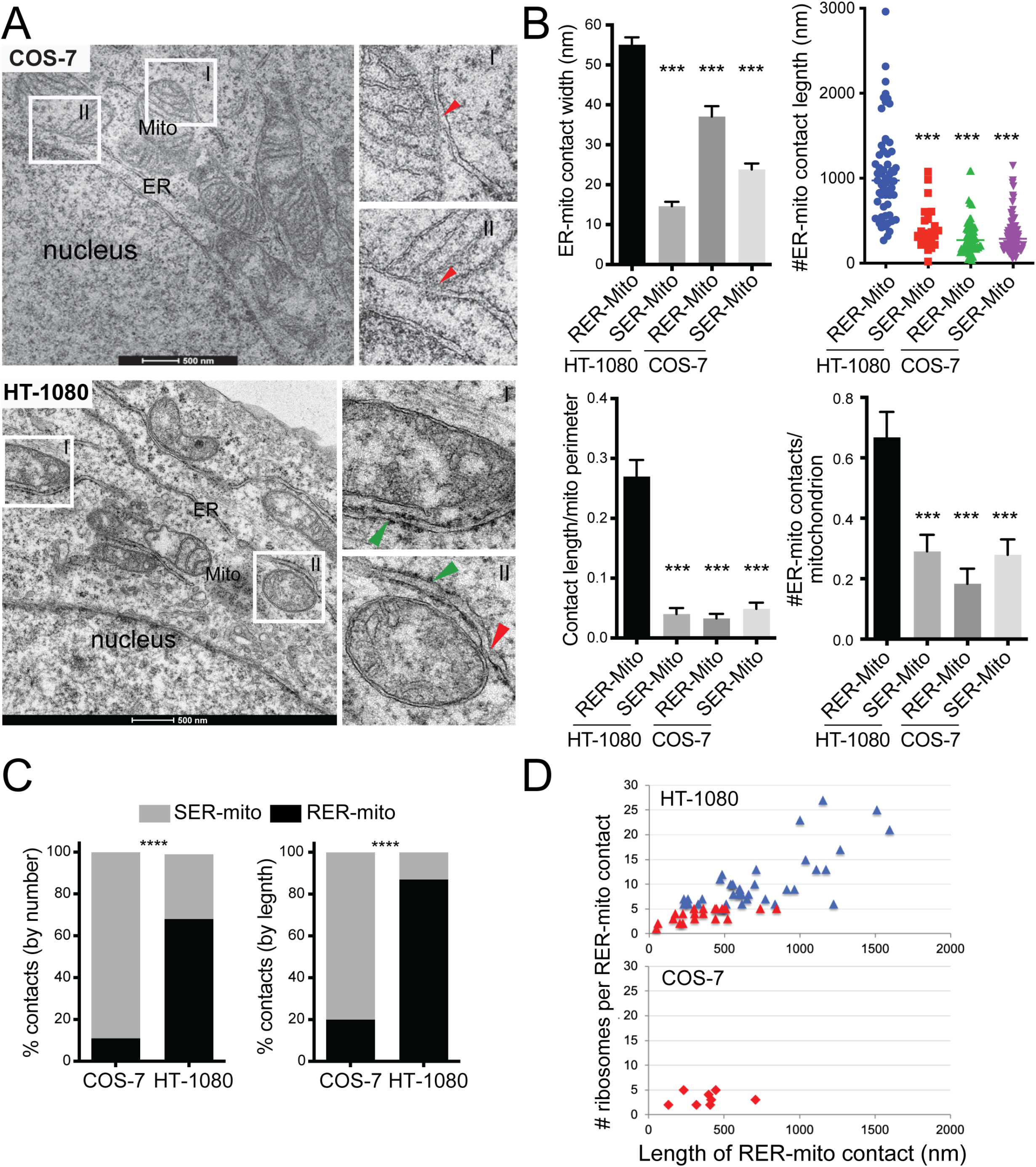
Quantitative EM analysis of ER-mitochondria contacts in HT-1080 and COS-7 cells. A) Representative EM images of HT-1080 and COS-7 cells. Insets show RER-mitochondria contacts (RER-mito) in HT-1080 cells and SER-mitochondria contacts (SER-Mito) in COS-7 cells. B) Quantification of contact width, contact length, contact length relative to mitochondria perimeter and number of contacts per mitochondria profile are shown for SER-mito and RER-mito contacts in HT1080 and COS-7 cells. C) The relative ratio of SER-mito and RER-mito contacts in HT-1080 and COS-7 cells based on number of contacts per mitochondria or length of contacts. D) The number of ribosomes per RER-mito contact is plotted vs length of the contact in nm for HT-1080 and COS-7 cells. RER-mito contacts with five or less ribosomes are shown in red; those with more than five ribosomes are specific to HT-1080 cells are shown in blue and defined as riboMERCs (+/-SEM; *** p < 0.001; **** p < 0.0001; Bar = 500nm).

RER-mito contacts in HT-1080 are of varying length and significantly longer than RER-mito contacts in COS-7 cells or SER-mito contacts in either HT-1080 or COS-7 cells (Fig. 1B). HT-1080 RER-mito contacts extend over almost 30% of the mitochondrial perimeter and contact a significantly larger proportion of mitochondria analyzed compared to SER-mito contacts or COS-7 RER-mito contacts that contact less than 5% of the mitochondrial surface. Almost 70% of HT-1080 MERCs are RER-mito contacts and compose 90% of MERCs in HT-1080 cells by length compared to less than 10% of MERCs in COS-7 cells (Fig. 1C). Importantly, RER-mito contacts of COS-7 cells contained at most 5 ribosomes within the interorganellar space compared to HT-1080 RER-mito contacts, some of which contained up to 25 ribosomes in a single contact (Fig. 1D). Elongated RER-mito contacts containing more than five ribosomes are therefore specific to HT-1080 cells and will heretofore be referred to as riboMERCs (Giacomello and Pellegrini, 2016).

Whole cell 3D STED image stacks of HT-1080 cells transfected with the lumenal ER reporter, ERmoxGFP (Costantini et al., 2015) were fixed with ER-preserving 3% paraformaldehyde/0.2% glutaraldehyde (Gao et al., 2019; Long et al., 2020; Nixon-Abell et al., 2016) and labeled for the outer mitochondrial membrane (OMM) protein TOM20 (Fig. 2A). Overlapping ER-mitochondria signal is observed in the cell periphery and more extensively in the central ER region, that for HT-1080 cells extends across multiple STED sections over 3 microns in depth. COS-7 cells present an abundance of ER-mitochondria overlapping regions but limited central ER overlap with mitochondria (Fig. 2A). The extended ER-mitochondria overlap of HT-1080 cells is consistent with the presence of elongated riboMERCs, as observed by EM (Fig. 1). However, defining actual contact sites from these 3D fluorescent whole cell views is challenging and subjective, based on user-dependent segmentation.

**Figure 2.**
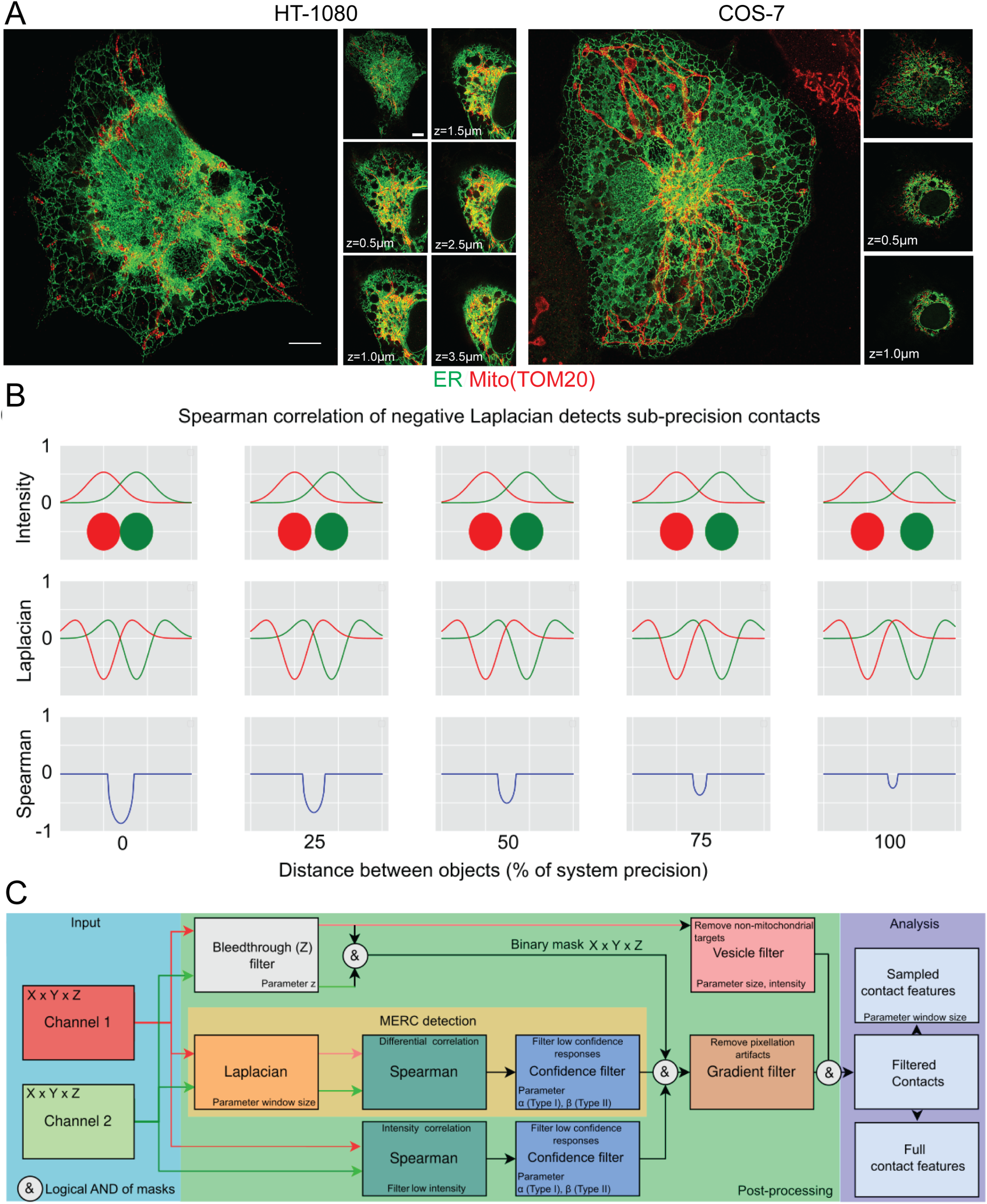
MCS-DETECT analysis of sub-precision contacts. A) 3D STED images of HT-1080 and COS-7 showing overlap between mitochondria (red) and ER (green). Insets show STED sections at 0.5 μM Z spacing. B) Two objects (red, green discs) are shown at corresponding sub-precision distances. Intensity profiles (top-row), 2^nd^ derivatives (Laplacian), and Spearman correlations of the negative part of the Laplacian (bottom row) are shown. Note how the Spearman response overlaps and changes consistently with the sub-precision distance. C) The detection algorithm (orange) with additional stages that each address a specific confounding factor introduced by the acquisition (bleedthrough) or sample (vesicle removal).

### MERC identification by differential channel correlation

Current image segmentation faces challenges of varying background density, difficulty of thresholding signals of varying density and particularly challenging when dealing with the highly varied density and complexity of the ER. In addition, analysis of overlap between two channels is based on separate segmentation of independent channels, compounding the error in capturing interaction. Here, we detect MERCs, below precision of the microscope, in this case 3D STED, by observing how the *relative* intensity of ER and mitochondria labeling change *in tandem*. Images are scanned using a sliding window applying a differential operator to approximate local signal differentials. Interaction between channels colocalizes with the negative Spearman correlation of the image differential (Fig. 2B). Contact zones are detected without requiring segmentation representing a novel approach to detect organelle contacts sites from multi-channel fluorescent images.

An in-silico experiment demonstrates the underlying principle of the method (Fig. 2B). An axial view of the intensity of two spheres acquired with a Gaussian point spread function (PSF) is shown in the upper row and negative Laplacian of the intensity profile of the two objects in the middle row. Distance between the 2 objects is varied from direct interaction up to the system resolution to mimic the MERC reconstruction problem where the interface is below acquisition precision. The detection principle relies on the negative correlation (bottom row, Fig. 2B) of the 2^nd^ intensity differential, approximated by the Laplacian operator. Importantly, the differential Spearman response is present for the entire sub-precision interaction range, confirming that a negative Spearman correlation of the negative Laplacian of the intensity profile corresponds to overlap of two adjacent objects, even when the precision of the system does not allow direct observation. The geometric mean of the Spearman correlation is reported for each contact. While the Spearman correlation can be impacted by variation in size and shape of the contact, at the cell level we observe consistent patterns across cell lines. As shown in Fig. 2C, the correlation principle is one step of a larger sequence of steps that address variable labelling density, low signal to noise ratio, intensity bleeding through multiple Z-slices, and anisotropic precision associated with 3D STED images. A full parameter sensitivity study is performed, with results shown in Supp. Fig. 1A. The full algorithm, including pseudocode and mathematical formulation for each stage, is detailed in Materials and Methods and Supp. Fig. 1B.

### Detecting riboMERCs in HT-1080 cells with MCS-DETECT

Application of MCS-DETECT to HT-1080 and COS-7 cells transfected with ERmoxGFP and labeled for mitochondrial TOM20 extracts a mask of contact zones (Fig. 3A). Overlay of the detected contacts on 3D volume rendering of deconvolved STED ER and mitochondria image stacks shows extended perinuclear contact zones in HT-1080 cells and smaller dispersed MERCs along mitochondria in COS-7 cells. Insets show that contact sites are localized to regions of interaction between ER (in green) and mitochondria (in red). 3D STED super-resolution microscopy is anisotropic with a predicted lateral resolution of 120 nm and axial resolution of 250 nm. Each voxel is 25×25×100 nm such that contact zone detection for larger contacts, and particularly for those that are located on top of mitochondria parallel to the plane of the acquired optical section, may be recovered with lower precision. We also detected a large number of contact sites between ER and smaller, reduced intensity TOM20-labeled mitochondrial structures (Supp Fig. 2), that may correspond to mitochondria vesicles (Neuspiel et al., 2008). To ensure that MERC analysis by MCS-DETECT parallels the contact sites detected by EM adjacent to intact mitochondria (Fig. 1), we filtered out these small, low intensity TOM20-labeled structures from the images based on size and intensity (Supp. Fig. 2).

**Figure 3.**
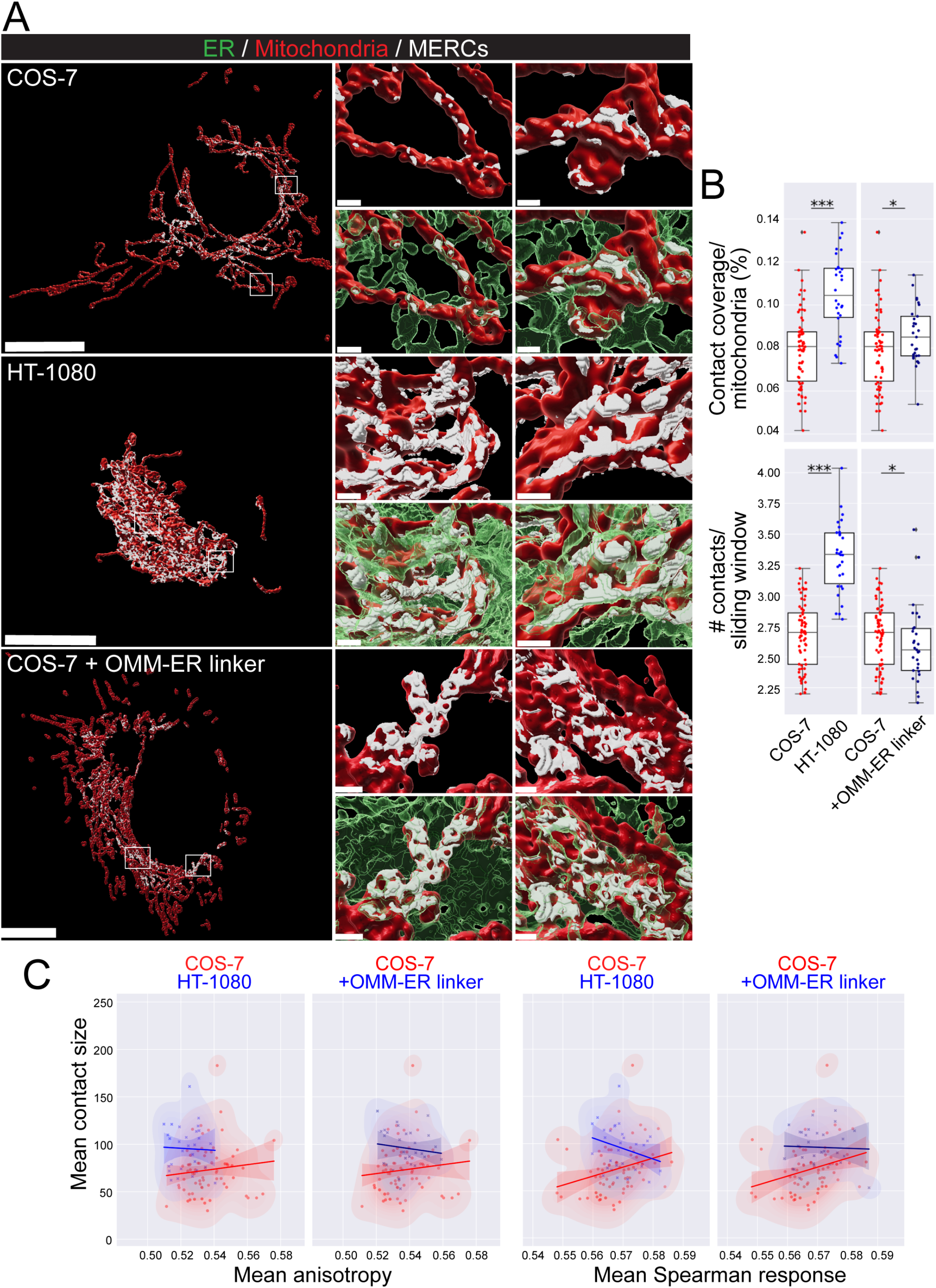
Sub-precision contact detection identifies distinct contact profiles in HT-1080 and COS-7 cells. A) Volume rendered MCS-DETECT views of cells expressing ERmoxGFP (green) and labelled for TOM20 (red) with contact sites overlaid (white) are shown for COS-7, HT-1080 and OMM-ER linker transfected COS-7 cells ROIs from the whole view image are shown volume rendered in adjacent panels. COS-7 mitochondria display numerous small contact zones while mitochondria in HT-1080 and OMM-ER linker transfected COS-7 cells present more extended contact zones. B) Mitochondria surface coverage ratio and the number of contacts per sampled mitochondria window are shown for contact zones in COS-7, HT-1080 and OMM-ER linker transfected COS-7 cells. C) 2D kernel density estimation (KDE) plots of mean contact size over mean anisotropy and mean spearman response, with a linear regression overlayed, are shown for COS-7 (red) vs HT-1080 (blue) cells or COS-7 (red) vs OMM-ER linker transfected COS-7 (blue) cells. (Averaged over cell, two-sided non-parametric Mann Whitney test, n=3; * p < 005; *** p < 0.001; Bar = 10 µm whole cell; 1 µm insets). 0.001; Bar = 10 µm whole cell; 1 µm insets).

To quantify ER-mitochondria contact zones in HT-1080 and COS-7 cells, we detected, per cell, mitochondria surface coverage ratio and number of mitochondria contacts within a fixed, non-overlapping, sliding window over the surface of each mitochondrion (Fig. 3B). HT-1080 cells present an approximate two-fold increased coverage of mitochondria and number of contacts per sample region, corresponding closely to the increased mitochondrial surface coverage we observe by EM (Fig. 1B), if one combines the smooth and riboMERC EM data as these are indistinguishable by STED microscopy. Pairwise analysis of MERC features shows that HT-1080 MERCs are on average larger and more spherical than COS-7 MERCs. A clear difference in anisotropy and mean Spearman response supports structural differences between MERCs of HT-1080 and COS-7 cells that show a clear correspondence with our EM results (Fig. 1).

Expression in COS-7 cells of a construct coding for the outer mitochondrial membrane AKAP sequence fused to the ER targeting sequence of Ubc, an OMM-ER linker, that induces close contacts (<20 nm) between ER and mitochondria (Csordas et al., 2010; Hajnoczky et al., 2006) induced extended contact sites (Fig. 3A). Mitochondrial surface coverage was increased relative to COS-7 cells but decreased relative to HT-1080 cells. In stark contrast to the MERCs of HT-1080 cells, a reduction in the number of contact sites per sample region was observed. As for HT-1080 cells, pairwise analysis showed that the MERCs of COS-7 cells expressing the OMM-ER linker were larger and showed distinct groupings compared to COS-7 MERCs with respect to both sphericity and Spearman response. The latter features were not however shifted away from COS-7 to the same extent as HT-1080 MERCs. The OMM-ER linker therefore induces expansion of COS-7 MERCs to form larger MERCs whose features are distinct from riboMERC expressing HT-1080 cells. MCS-DETECT therefore detects MERC features that form a statistically significant signature for diverse type of contacts.

### Gp78 and RRBP1 are independent regulators of riboMERC expression

Using CRISPR/Cas knockout Gp78 HT-1080 cells that present deficient basal mitophagy, impaired mitochondrial health and increased mitochondrial ROS (Alan et al., 2022), we show that loss of Gp78 reduces MERC mitochondrial coverage (Fig. 4A). Surface coverage and number of contacts per mitochondrial surface region are significantly reduced relative to HT-1080 cells but remain elevated compared to COS-7 cells (Fig. 4B). The partial reduction of riboMERCs upon Gp78 KO is consistent with prior EM results showing that Gp78 knockdown does not completely eliminate riboMERCs (Wang et al., 2015). Overexpression of Gp78 in COS-7 cells induces the inverse response in which COS-7 cells gain large contacts similar to those found in HT-1080 cells. Importantly, overexpression of a RING finger mutant of Gp78, Gp78 RM, lacking ubiquitin ligase activity and unable to induce mitophagy (Fang et al., 2001; Fu et al., 2013), is unable to induce this effect. Shape feature analysis shows that Gp78 expression in COS-7 cells induces size-dependent increase in anisotropy and reduction of mean Spearman response that match those observed in HT-1080 cells (Fig. 4C). These data indicate that the large contact zones selectively enriched in HT-1080 cells correspond to riboMERCs and identify Gp78 ubiquitin ligase activity as a specific regulator of the size of these contact sites.

**Figure 4.**
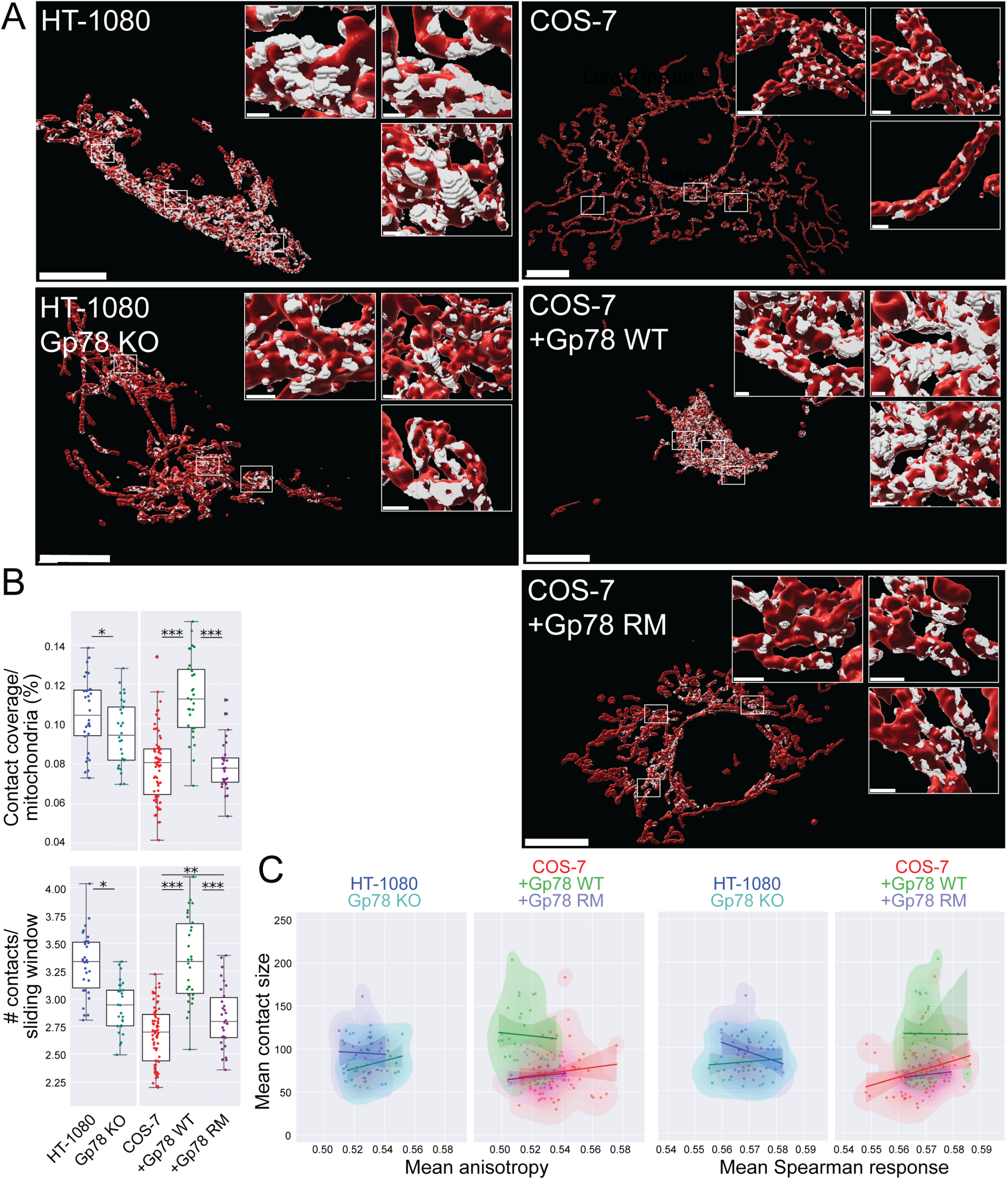
Gp78 regulation of riboMERCs. A) Volume rendered MCS-DETECT views of cells expressing ERmoxGFP and labelled for TOM20 (red) with contact sites overlaid (white) are shown for HT-1080 and Gp78 KO HT-1080 cells and for untransfected COS-7 cells and COS-7 cells overexpressing wild-type (WT) Gp78 or RING-finger domain mutated Gp78 (Gp78 RM). B) Mitochondria surface coverage ratio and the number of contacts per sampled mitochondria window are shown for contact zones in HT-1080 and Gp78 KO HT-1080 cells and for untransfected COS-7 cells and COS-7 cells overexpressing Gp78 WT or Gp78 RM. C) 2D KDE plots of mean contact size over mean anisotropy and mean spearman response, with a linear regression overlayed, are shown for HT-1080 (blue) vs Gp78 KO HT-1080 (green) cells or COS-7 (red) vs COS-7 overexpressing either Gp78 WT (green) or Gp78 RM (blue). (Averaged over cell, two-sided non-parametric Mann Whitney test, n=3; * p < 0.05; ** p < 0.01; *** p < 0.001; Bar = 10 µm whole cell; 1 µm insets).

Previous studies identified RRBP1 as a MERC resident protein required for increased riboMERC expression through interaction with its mitochondrial partner SYNJ2BP (Hung et al., 2017). To investigate the impact of RRBP1 on riboMERC formation in HT-1080 cells, wild-type and Gp78 KO HT-1080 cells were treated with either control siRNA or siRNA targeted to RRBP1 and analyzed by EM (Fig. 5). Wild-type HT-1080 cells displayed both the highest number of riboMERCs per mitochondria as well as the highest ratio of riboMERC length to mitochondrial perimeter which showed a partial reduction in Gp78 KO cells, consistent with our MCS-DETECT analysis (Figure 4) and prior EM studies of Gp78 shRNA knockdown HT-1080 cells (Wang et al., 2015). However, upon treatment with siRRBP1, we observed an almost complete ablation of riboMERCs (Fig. 5B). Analysis of MERC width revealed that the few remaining riboMERCs in siRRBP1 treated Gp78 KO cells had a slight if significantly larger width (Fig. 5C), perhaps reflecting increased spacing of riboMERCs as observed in siRRBP1-treated hepatocytes (Anastasia et al., 2021). MCS-DETECT analysis of 3D STED super-resolution images of siRRBP1 transfected cells mirrored the EM analysis; RRBP1 knockdown decreased both the absolute number of contacts as well as percentage of the mitochondrial surface covered by MERCs, irrespective of Gp78 expression (Fig. 6A, B). These values in siRRPB1 treated HT-1080 cells were similar to those observed in COS-7 cells (Figure 4), that also show an absence of riboMERCs. Similarities in MCS-DETECT reporting on MERC coverage in two cell lines, COS-7 in which riboMERCs are induced by Gp78 expression and HT-1080 in which they are lost by siRRBP1 knockdown, highlights the robustness of the MERC detection approach.

**Figure 5.**
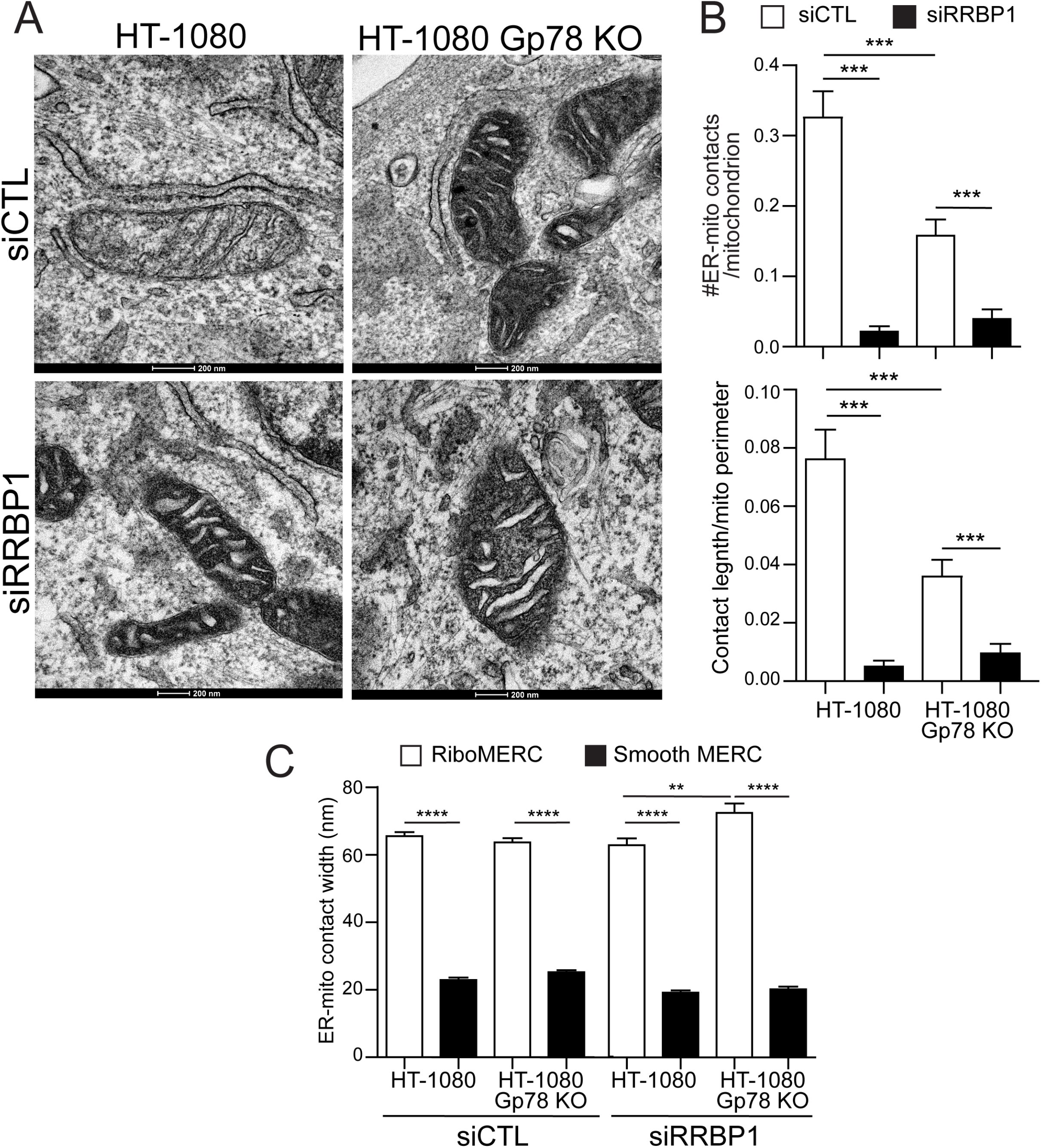
RRBP1 knockdown reduces riboMERCS independent of Gp78. A) Representative EM images of HT-1080 and HT-1080 Gp78 KO cells treated with either siControl or siRRBP1. Images highlight the presence of riboMERCS in both HT-1080 WT and Gp78 KO cells which are almost completely lost upon RRBP1 knockdown. B) Quantification of the number of riboMERCs per mitochondria and the ratio of riboMERC length to mitochondrial perimeter for the conditions in A. C) Quantification of the MERC width for both riboMERCs and smooth MERCs for the conditions in A.

**Figure 6.**
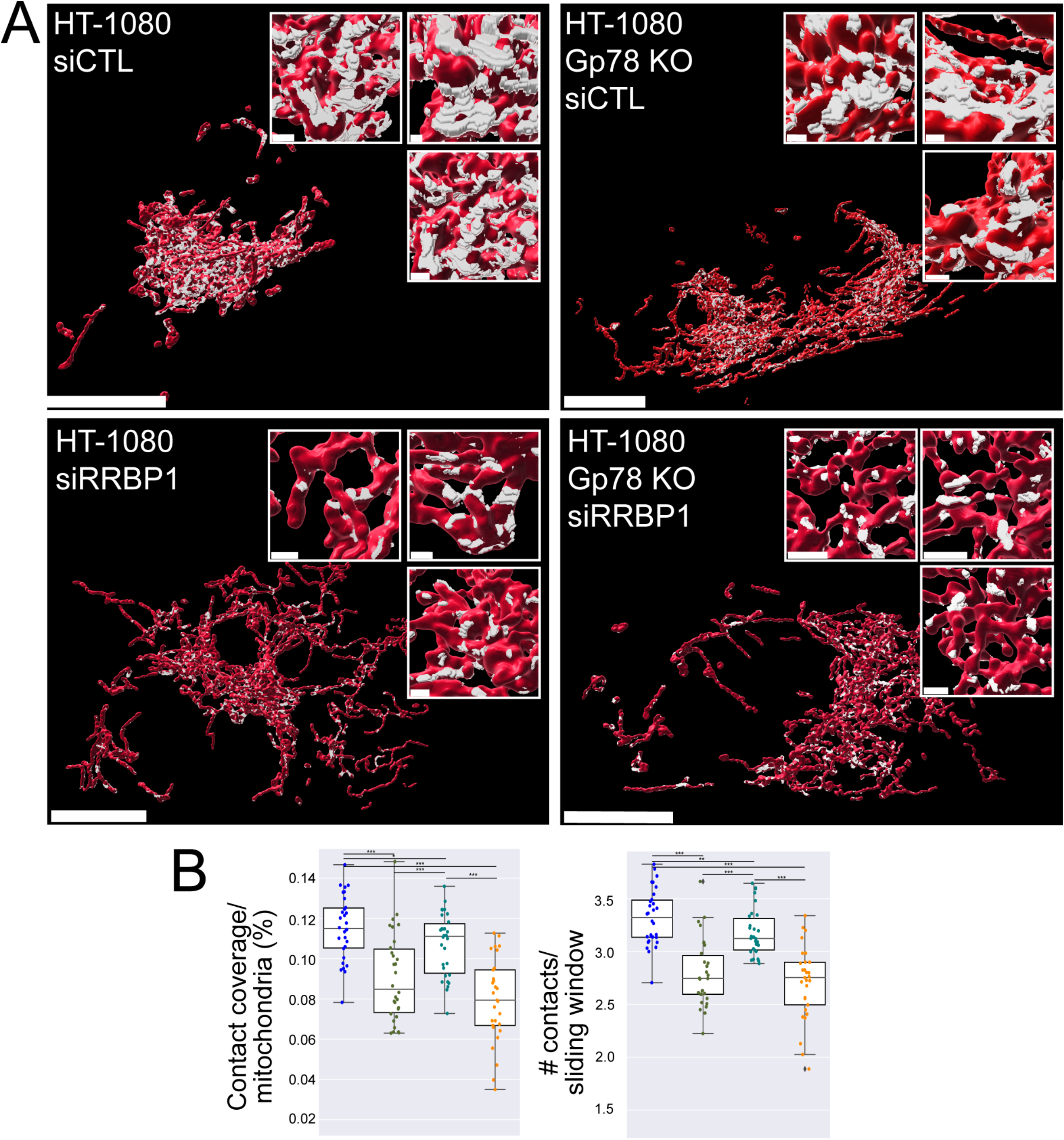
MCS-DETECT captures MERC changes induced by RRBP1 knockdown. A) Volume rendered MCS-DETECT views of HT-1080 WT and Gp78 KO cells treated with either siControl or siRRBP1. Mitochondria are labelled with TOMM20 (red) and MERCS are visualized in white. B) Mitochondria surface coverage ratio and the number of contacts per sampled mitochondria window are shown for contact zones in HT-1080 WT and Gp78 KO cells treated with either siControl or siRRBP1. C) 2D KDE plots of mean contact size over mean anisotropy and mean spearman response, with a linear regression overlayed, are shown for HT-1080 (blue) vs HT-1080 siRRBP1 (green) cells or HT1080 Gp78 KO cells (Cyan) vs Gp78 KO cells treated with siRRBP1 (Orange). (Averaged over cell, two-sided non-parametric Mann Whitney test, n=3; * p < 0.05; ** p < 0.01; *** p < 0.001; Bar = 10 µm whole cell; 1 µm insets).

### Tubular, Gp78-dependent riboMERCs

To characterize riboMERCs at the level of individual MERCs, we analyzed the largest 5% of MERCs in each cell, which based on our EM analysis correspond to elongated riboMERCs in HT-1080 cells (Fig. 1). Analysis of the size of the largest 5% of MERCs per cell (Volume Q95) in the various cells and treatments analyzed (Fig. 5A) paralleled our analysis of the average size of all MERCs per cell (Fig. 3B, 4B). This represents a strong indication that differences in contacts across cell lines are driven by changes in the tail (largest) of the contact size distribution. The highly significant differences between the largest 5% MERCs between HT-1080 and both COS-7 and siRRBP1 treated HT-1080 cells indicates that the largest 5% of HT-1080 MERCs encompasses predominantly riboMERCs. Overexpression of both Gp78 and the OMM-ER linker in COS-7 cells increased the size of top 5% MERCs, with Gp78 induced Q95 MERCs even larger than those of wild-type HT-1080 cells. We further counted the number of large MERCs, larger than the 500 voxel average size of the largest 5% of HT-1080 MERCs (Figure 7A). The number of large MERCs was significantly reduced in COS-7 relative to HT-1080 cells and upon siRRBP1 knockdown in HT-1080, consistent with the absence of riboMERCs in those cells (Figs. 1, 5). The number of large MERCs was increased in COS-7 cells upon expression of the OMM-ER-linker and even more so upon expression of Gp78. Interestingly, the top 5% of MERCs in Gp78 KO HT-1080 cells were smaller than in HT-1080 cells yet the number of large (>500 voxel) MERCs was the same in the two cell lines suggesting that Gp78 KO selectively regulates the size of riboMERCs as opposed to their abundance. Together, these EM and MCS-DETECT data demonstrate that, in contrast to RRBP1 that is essential for riboMERC expression in HT-1080 cells, Gp78 rather acts to modulate the size of riboMERCs via its ubiquitin ligase activity.

**Figure 7.**
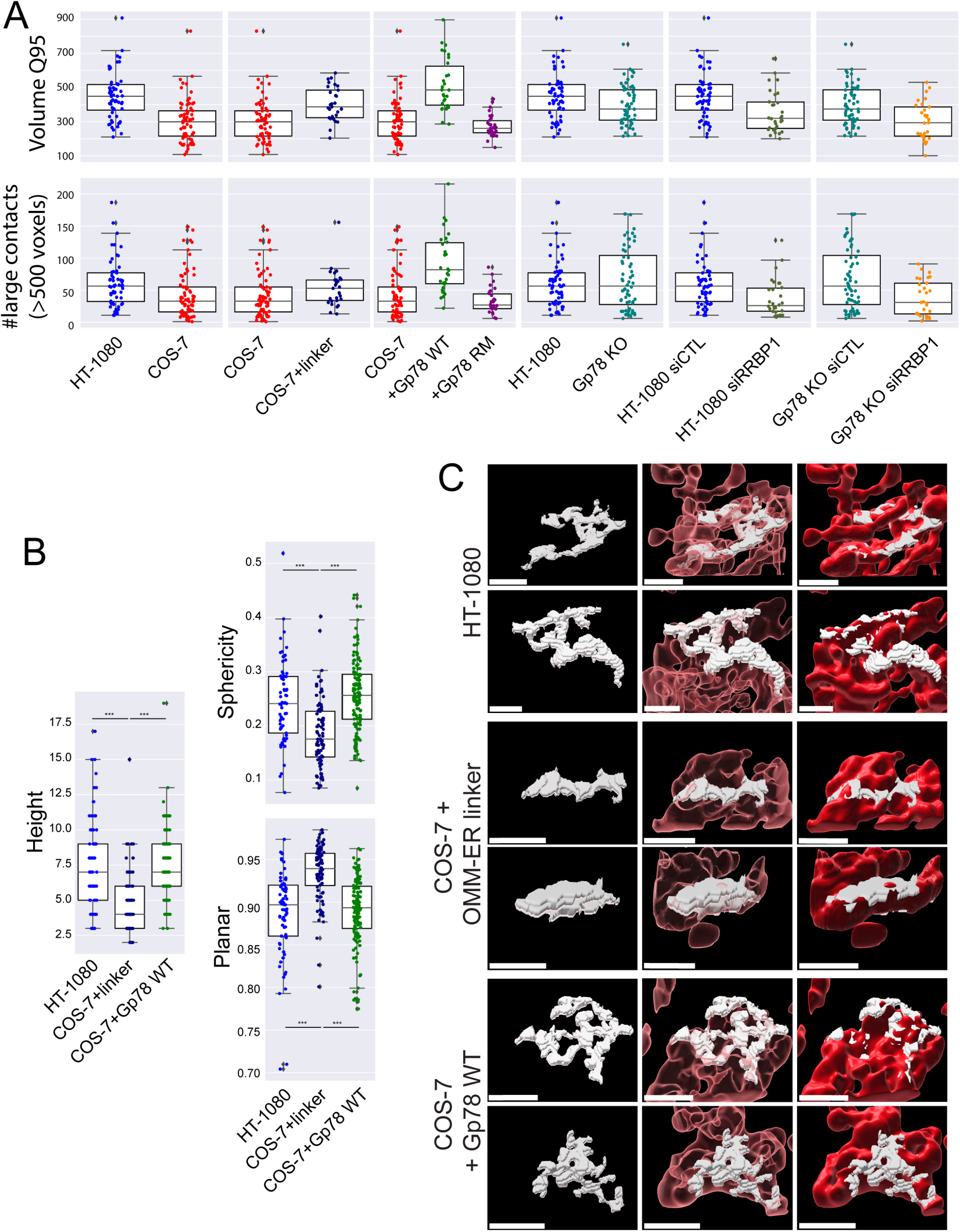
Gp78 induces convoluted, tubular riboMERCs. A) The 95^th^ quantile of MERC volume per cell (Q95V; largest 5% of MERCs per cell) and number of MERCs per cell larger than the average 500 voxel size of HT-1080 Q95V MERCs are shown for HT-1080 (blue) and COS-7 (red) cells, COS-7 (red) and COS-7 overexpressing the OMM-ER linker (black), COS-7 (red) and COS-7 overexpressing either Gp78 WT (green) or Gp78 RM(purple), HT-1080 (blue) and Gp78 KO HT-1080 (teal) cells, HT-1080 cells transfected with siCTL (blue) and siRRBP1 (sage green), and Gp78 KO HT-1080 cells transfected with siCTL (teal) and siRRBP1 (orange). B) A representative cell whose Q95V is closest to the mean Q95V of the HT-1080 (blue) cells and COS-7 cells overexpressing either the OMM-ER linker (black) or Gp78 WT (green) was selected for analysis. For the Q95V contacts of each cell we compute shape features: height, sphericity and planarity. The comparison shows that the COS-7 OMM-ER linker induced contacts have a markedly different shape signature compared to those present in HT-1080 and COS-7 Gp78OE (i.e. riboMERCS). C) Representative views of Q95V MERCs (white) from HT-1080 cells and COS-7 cells expressing either the OMM-ER linker or wild-type Gp78 are shown alone or adjacent to transparent (pink) or solid mitochondria (red) to highlight intercalation of riboMERCs with mitochondria. Rotating videos of these MERCs are attached as Supplemental Videos. Bar = 1 μm.

Shape feature analysis of the largest 5% MERCs shows that riboMERCs of HT-1080 cells are taller, more spherical and less planar than the MERCs induced by the OMM-ER linker in COS-7 cells (Fig. 7B). RiboMERCs of HT-1080 cells form extended, tubular structures that are intercalated between and form contacts with multiple mitochondria (Fig. 7C). The large MERCs induced by the OMM-ER linker form more elongated planar structures that are closely associated with an individual mitochondrion, similar to the cap-like structures observed by EM upon expression of this linker (Csordas et al., 2006). The different morphology of OMM-ER linker induced MERCs explains the decrease in number of contact sites per region observed upon expression of the OMM-ER linker in COS-7 cells (Fig. 3B). Features and structure of large MERCs induced by overexpression of Gp78 in COS-7 cells are similar to that of HT-1080 riboMERCs, demonstrating that Gp78 induces the formation of extended, tubular riboMERCs in COS-7 cells (Fig. 7C). MCS-DETECT is therefore able to distinguish MERCs based on size and shape features defining the riboMERCs of HT-1080 cells as extended, tubular matrix like structures that interact with and intercalate between multiple mitochondria. Detection of riboMERC structural changes by MCS-DETECT highlights the importance of whole cell 3D analysis of MERCs, as these changes could not be detected in our 2D EM analysis, illustrating the potential for discovery MCS-DETECT brings to the study of MERCs.

## DISCUSSION

MERCs were initially characterized based on biochemical identification of mitochondria-associated membranes (MAM), that identified a role for MERCs in lipid synthesis (Vance, 1990; Vance, 1991). MERCs were subsequently shown to play critical roles in ER-mitochondria calcium transport, cellular calcium homeostasis and apoptosis (Csordás et al., 2018; Herrera-Cruz and Simmen, 2017; Rowland and Voeltz, 2012). However, while use of functional reporters and cell fractionation approaches to define ER-mitochondria contacts led to important understanding of the role and composition of MERCs, these approaches are unable to localize MERCs within the cell (Scorrano et al., 2019). Reliance on EM to detect MERCs and associated difficulties in localizing tethers to MERCs has led to discordant results as to whether specific proteins, for instance MFN2, are indeed MERC tethers (Cosson et al., 2012; de Brito and Scorrano, 2008; Dentoni et al., 2022; Leal et al., 2016; Naon et al., 2016; Wang et al., 2015).

Moving beyond EM to use fluorescent microscopy to characterize MERC composition and dynamics includes the development of split fluorescent reporters (such as SPLICs (Cali and Brini, 2021; Cieri et al., 2018; Vallese et al., 2020) whose expression may however alter MERC size and stability. The use of fluorescent colocalization to detect MERCs is particularly problematic as the distance between the ER and mitochondria at MERCs (10-80 nm) is far below the diffraction-limit (200-250 nm) of visible light used for fluorescent microscopy. Further, the poorer axial resolution of most fluorescent microscopy approaches, including super-resolution, have restricted fluorescent microscopy analysis of MERCs to peripheral ER tubules (Friedman et al., 2011) such that analysis of mitochondria interaction within the convoluted perinuclear ER remains poorly understood. Here, using STED super resolution imaging and a novel segmentation-free sub-pixel resolution approach to identify membrane contact sites (MCS-DETECT), we identify MERCs in whole cell 3D volumes of HT-1080 and COS-7 cells and further detail the role of Gp78 ubiquitin ligase activity and the riboMERC tether RRBP1 in the abundance and morphology of riboMERCs.

Current image segmentation using fixed thresholds faces challenges of varying background density, difficulty of thresholding signals of varying density and is highly subjective and user-dependent. In addition, analysis of overlap between two channels is based on separate segmentation of independent channels, such that error in capturing interaction, in this case MERCs, is unknown. An optimal method adapts automatically to the data, ensuring consistent results, across images and datasets. To this end we developed a multichannel self-tuning object detection method (SPECHT), building upon our work in SMLM object detection (ERGO (Cardoen et al., 2022; Cardoen et al., 2020). To enhance object (spot) detection, SPECHT scans the image and applies a Laplacian differential operator to detect local signal differentials and identify object edges. Automated image scanning (a sliding window) and kurtosis-based image alignment results in highly sensitive detection independent of variations in local signal or background intensity differentials. This approach is optimal to segment the ER, an organelle of highly varied density and complexity. To validate our object detection method, we generate complex in silico datasets to accurately detect interaction of objects in low signal to noise ratio (SNR) channels, especially where the SNR varies markedly between channels. To identify interaction zones between two channels, we identify overlapping negative intensity differentials of both channels (mitochondria, ER); overlap is only detected where edges of the two signals are in sub-resolution space (i.e. lower than the 3D STED resolution of ∼100 nm). Laplacian operators are dependent on local signal differential for each channel such that detection of overlapping regions is independent of signal intensity. The Spearman correlation of overlapping ER and mitochondrial Laplacians defines MERC regions that are described by volume and voxel-wise Spearman response. MCS-DETECT therefore represents a novel sub-pixel resolution approach ideally suited to detect contact zones between organelles.

Morphologically distinct riboMERCs are wider, include ribosomes in the intervening space and are abundantly expressed in liver (Anastasia et al., 2021; Csordas et al., 2006). They are less abundant in most cultured cells, as shown here in our analysis of COS-7 cells, but were found to be abundant by EM analysis of HT-1080 fibrosarcoma cells, that express robust amounts of the Gp78 ubiquitin ligase (Alan et al., 2022; Tsai et al., 2007). Gp78 knockdown in HT-1080 cells selectively reduced riboMERC expression (Wang et al., 2015), a result we confirm here using CRISPR/Cas Gp78 KO HT-1080 cells. The demonstration here that wild-type Gp78 but not a Ring finger mutant Gp78 can induce large riboMERCs in COS-7 cells defines a role for Gp78 ubiquitin ligase activity in riboMERC formation. Gp78 regulation of riboMERCs may involve interaction with ubiquitinated mitochondrial partners, such as the mitofusins (Fu et al., 2013; Mukherjee and Chakrabarti, 2016). However, the presence of riboMERCs in Gp78 KO cells clearly shows that other factors promote the formation of riboMERCs independently of Gp78. Interaction between the OMM protein SYNJ2BP/OMP25 and its ER partner, ribosome-binding protein 1 (RRBP1) was shown to induce the formation of riboMERCs in HEK293 cells (Hung et al., 2017). Knockdown of RRBP1 results in reduced riboMERC formation and increased spacing of riboMERCs in liver (Anastasia et al., 2021). Our demonstration that RRBP1 knockdown in HT-1080 eliminates riboMERCs defines an essential role for RRBP1 in the formation of riboMERCs. RRBP1 tethering activity, through interaction with its mitochondrial partner SYNJ2BP (Hung et al., 2017), occurs independently of Gp78. Regulation of RRBP1-dependent riboMERCs by Gp78 ubiquitin ligase activity therefore represents a novel paradigm for MERC formation, in which an effector protein modulates the extent to which a MERC tether can form contact sites.

3D EM tomography studies of riboMERCs in liver show that they that wrap completely around mitochondria, covering 30-100% of associated mitochondria, and were therefore labeled WrappER (Anastasia et al., 2021). In contrast, riboMERCs in HT-1080 cells do not wrap completely around mitochondria. By thin section EM (Fig. 1), riboMERCs form extended contacts along a single face of the mitochondrion that cover at most 30% of the mitochondrion perimeter (Figure 1). While riboMERCs in liver present a sheet-like morphology (WrappER) that wraps around the mitochondrion (Anastasia et al., 2021), 3D STED imaging shows that Gp78-dependent riboMERCs form convoluted, tubular structures resembling tubular matrices (Nixon-Abell et al., 2016) that intercalate between and interact with multiple mitochondria. Consistent with a tubular morphology, the MERCs of HT-1080 and Gp78-transfected COS-7 cells show increased contact sites per analysis window relative to COS-7 cells. In contrast to the tubular Gp78-dependent riboMERCs, MERCs induced by the OMM-ER linker formed more planar structures. The decrease in contact number for OMM-ER linker expressing COS-7 cells is highly consistent with the tight (<20 nm) extended planar contacts over the surface of a mitochondrion formed upon expression of this interorganellar linker (Csordas et al., 2006). An increased number of contacts per sliding window could reflect varied distance between ER and mitochondria within the contact, as observed for liver riboMERCs upon knockdown of the riboMERC tether RRBP1 (Anastasia et al., 2021; Hung et al., 2017). Contact width of HT-1080 riboMERCs is very constant ranging from 50-60 nm along the length of the contact (Wang et al., 2015). We interpret this to mean that HT-1080 riboMERCs form extended tubular networks that form multiple contacts with mitochondria.

As for tubular matrix ER and ER sheets (Nixon-Abell et al., 2016; Sun et al., 2020), the functional and morphological relationship between tubular riboMERCs and sheet-like WrappER remains to be determined. WrappER is implicated in lipid homeostasis and recently shown to interact with peroxisomes in liver (Anastasia et al., 2021; Ilacqua et al., 2022) while Gp78 ubiquitin ligase activity in HT-1080 cells mediates basal mitophagy, promoting mitochondrial health and reducing mitochondrial ROS (Alan et al., 2022). Further definition of these and, potentially, other functions for riboMERCs requires further study. Indeed, this study highlights the fact that MERCs can take on diverse morphologies. While this analysis focused on a select subset of riboMERCs and the riboMERC regulators Gp78 and RRBP1, the large number of MERC tethers identified suggests a high degree of diversity of MERCs, with respect to both function and structure (Giacomello and Pellegrini, 2016; Herrera-Cruz and Simmen, 2017).

Study of the diversity of MERCs, their associated tethers and specific functionality has been limited by the absence of a robust fluorescence-based approach to detect MERCs. Application of MCS-DETECT to 3D super-resolved volumes of fluorescently labeled cells now provides a means to extend our characterization of MERC diversity, localize MERC tethers to contact sites and study MERC dynamics, complementing other criteria used to define membrane contact sites (Scorrano et al., 2019). Importantly, MCS-DETECT is not dependent on expression of MERC reporter molecules and, relative to EM tomography or FIB-SEM, able to rapidly analyze multiple cell volumes. While we used here 3D STED fixed cell analysis of established models for riboMERC expression to validate the approach, MCS-DETECT reports on negative Laplacians of any two overlapping fluorescent signals and can easily be applied to contacts between any two organelles detected with any voxel-based super-resolution system.

## MATERIALS AND METHODS

### Cell culture

COS-7, HT-1080 and Gp78 KO HT-1080 (Alan et al., 2022) cells were grown at 37°C with 5% CO_2_ in complete DMEM or RPMI 1640 (HT1080 WT and Gp78 KO) or DMEM (COS7) (Thermo Fisher Scientific, USA) containing 10% FBS (Thermo Fisher Scientific, USA) and 1% L-Glutamine (Thermo Fisher Scientific, USA) unless otherwise stated. Plasmids were transfected using Effectene (Qiagen, Germany) according to the manufacturer’s protocols for 22 hours. siRNA was transfected using Lipofectamine 2000 (Thermo Fisher Scientific, USA Cat#: 11668019) according to the manufacturer’s recommendation for 48 hours.

### Antibodies, plasmids, and chemicals

ERmoxGFP was a gift from Erik Snapp (Addgene plasmid # 68072). Goat serum was purchased from Thermo Fisher Scientific (Cat#: 16210-064), mouse monoclonal (Cat#: ab56783) to TOM20/TOMM20 (1:300 dilution) from Abcam, 16% paraformaldehyde (Cat#: 15710), and 25% glutaraldehyde (Cat#: 16220) were from Electron Microscopy Sciences (Hatfield, PA, USA). RRBP1-targeted and scrambled control siRNA were purchased from Dharmacon (Cat#: L-011891-02-0010). Other chemicals were from Sigma.

### Electron microscopy

Cells grown on ACLAR film (Ted Pella) for 24 hours until 80-90% confluency, where indicated transfected with specific siRNAs for an additional 48 hours, were (1) washed with PBS at room temperature, (2) fixed with 1.5% paraformaldehyde and 1.5% glutaraldehyde in 0.1M sodium cacodylate (Electron Microscopy Sciences, Hatfield, PA, USA), pH 7.3 for 1 h at room temperature followed by 3 washes with 0.1M sodium cacodylate, pH 7.3, (3) postfixed with 1% osmium tetroxide in 0.1M sodium cacodylate buffer for 1 h at 4°C on ice following by 3 washes with H_2_O, (4) stained *en bloc* with aqueous 1.0% uranyl acetate for 1 h on ice, (5) progressively dehydrated through an ethanol series (50%, 70%, 90%, and 100%) and 100% propylene oxide (Electron Microscopy Sciences, Hatfield, PA, USA) each for 10 min, (6) infiltrated with 1:1 mixture of EMBED 812 (Electron Microscopy Sciences, Hatfield, PA, USA) and propylene oxide, and embedded in EMBED 812, (7) polymerized for 2 days, (8) sectioned into approximately 900 A thick slices using Leica Ultramicrotome (Leica Microsystems, Wetzlar, Germany), (9) stained with uranyl acetate and lead citrate, (10) imaged either on a Tecnai G2 (FEI) or a Talos L120C (ThermoFisher) transmission electron microscope (acceleration voltage: 120 kV), using an Eagle 4k CCD camera (FEI) or a CETA camera (ThermoFisher), respectively.

### Immunofluorescence labeling

Cells grown on #1.5H coverslips (Paul Marienfeld, Germany) were: 1) fixed with 3% paraformaldehyde with 0.2% glutaraldehyde at room temperature for 15 minutes and washed with PBS-CM (phosphate buffer solution supplemented with 1mM CaCl_2_ and 10mM MgCl_2_) (two quick washes and then two 5 minute washes); 2) permeabilized with 0.2% Triton X-100 for 5 minutes then washed with PBS-CM as above; 3) quenched with 1mg/mL of NaBH_4_ for 10 minutes and washed with PBS-CM; 4) blocked with 10% Goat Serum (Thermo Fisher Scientific, USA) and 1% bovine serum albumin (Sigma, USA) in PBS-CM for 1 hour; 5) incubated with primary antibodies in Antibody Buffer (1% BSA, 2% goat serum, 0.05% Triton-×100, 20X sodium/sodium citrate buffer in Milli-Q H_2_O) overnight at 4°C then washed quickly with PBS-CM then three times for 5 minutes with Antibody wash buffer (20x SSC, 0.05% Triton-×100 in Milli-Q H_2_O); 6) incubated with secondary antibodies in Antibody Buffer for 1 h then washed quickly with PBS-CM then six times for 10 minutes with Antibody Wash Buffer on a rocker; 7) rinsed with Milli-Q H_2_O and mounted with ProLong Diamond (Thermo Fisher Scientific, USA) and cured for 24-48 hours at room temperature.

### gSTED microscopy

gSTED imaging were performed with the 100X/1.4 Oil HC PL APO CS2 objective of a Leica TCS SP8 3X STED microscope (Leica, Germany) using white light laser excitation, HyD detectors, and Leica Application Suite X (LAS X) software. Time-gated fluorescence detection was used for STED to further improve lateral resolution. For double labeled fixed samples, the acquisition was done at a scan speed of 600Hz with a line average of 3. GFP was excited at 488 nm and depleted using the 592 nm depletion laser. Alex Fluor 568 was excited at 577 nm and depleted using the 660nm depletion laser. Sequential acquisition (in the orders of AF568/GFP) between frames/stacks was used to avoid crosstalk. STED images were deconvolved using Huygens Professional software (Scientific Volume Imaging, Netherlands).

### MCS-DETECT: Contact zone detection pipeline

The source code is available under an open source license (AGPLv3) at https://github.com/bencardoen/SubPrecisionContactDetection.jl. An optimized version with all dependencies in a single container image is provided as well for ease of use.

Illustrated in Figure 2C, the algorithm is composed of 3 steps: computing correlation of intensity and Laplacian for both channels, computing the confidence map, and finally filtering to remove artifacts. A full listing in pseudocode of each stage is provided in Supp. Fig. 1B.

### Spearman correlation

We retain the negative Spearman response on the negative Laplacian of each channel (Supp. Fig 1B Alg. 1, 3). In order to prevent the inclusion of low intensity signals colocalizing with high intensity signals, we compute the negative Spearman response of the intensity of both channels; only where both Laplacian and intensity correlate, do we retain the Laplacian correlation.

### Confidence map

For each Spearman response voxel, we compute its significance and the minimum observable correlation (Supp. Fig. 1B; Alg. 4, 5), given the sample size (window), using a z-test. Typical values used here are 0.05 both for alpha and statistical power (1-beta). In Supp. Fig. 1A we show that this parameter leads to consistent results on representative cells.

### Filtering

Because 3D STED microscopy is often anisotropic in its point spread function, and one thus risks ‘bleed through’ across z-planes, we apply a ‘bleedthrough’ filter (Supp. Fig. 1B; Alg. 2) as a mask by filtering out intensity lower than a given z-score (standard score of intensity), where z is the parameter set in the filter. Bleed-through intensity values will have markedly smaller z-values compared to ER and mitochondria segments. Because the z-score is a pivotal quantity, it will adapt to each channel’s distribution and thus lead to an unbiased filter. We test the effect of this filter in Supp. Fig. 1B, where consistent results are retained across a large range of the parameter. Due to low SNR and pixelation, a ‘shadowing’ response, parallel at an offset of 3-4 voxels to a true response, can appear. As can be observed from the Laplacian curves in Fig 2-B (middle row), voxels where only Laplacian of only one channel changes cannot colocate with a contact. Our gradient filter computes the change of the Laplacian by way of the 3^rd^ derivative of intensity. Next, we mask out any voxels where this 3^rd^ derivative is 0 for 1 channel, yet nonzero for the other. The interested reader can infer from Fig. 2B that a 3^rd^ derivative can be zero for a valid contact, but only if both are zero.

To ensure that we report on contacts with whole mitochondria and not mitochondria fragments or vesicles, as we do by EM, we exclude contacts that are adjacent to mitochondria below a size and mean intensity threshold of respectively 9 (ln scale, voxels) and 0.2, chosen based on empirical observation of non-target mitochondria-labeled segments. We illustrate that these thresholds split the distribution of mitochondria between mitochondria and vesicle-like objects in Supp. Fig. 2. For both COS-7 and HT1080 cells a clear bimodal distribution of mitochondria segments is separable.

### Quantifying Contacts

We compute features of contacts sampled by both a sliding window and per-contact. Contacts are distributed with a frequency strongly inverse to their size. Quantifying such long tail distributions can undercount the visually more apparent large contacts. To this end we slide a non-overlapping window over the contact channel and record size and frequency per cube, estimating local density. At the whole contact level, we record the confidence, shape, size, and spatial distribution of the contacts. The resulting 2 sets of features allow the end user a local and global descriptive view of contacts in the cell.

For our data we use a sliding window of 25×25×5 voxels, which equates to a cube given the anisotropy of the acquisition. We show in the Supp. Fig. 2A how this window size does not alter the consistency of our results. We average results over cells and test the result with a non-parametric two-sided Mann Whitney test. Applying multiple testing correction is not practical here, given that we have a pairwise (HT-1080), and 3 pairwise comparisons (COS-7), meaning the correction would be applied only to COS-7 we would be reporting results of COS-7 with a risk of false negative higher than for HT-1080. Contacts are analyzed both by sampling the image, but also as a whole. Large contacts are defined as those exceeding the 95^th^ quantile of volume, per cell. Contacts with size <= 2 voxels are removed, as they are below the diffraction limit. We compute shape features (sphericity, anisotropy, planarity) based on the eigenvalues of the Spearman weighted contacts. Not reported but provided to users of MCS-DETECT are: confidence of reconstruction, clustering, position in Z, to name a few.

### Statistical Analysis

Statistical tests are always computed on aggregated (per cell) results to avoid breaking the independence assumption or inducing Simpson’s paradox. No outlier removal is performed. Plots are edited for cosmetic alterations only. To avoid dependence on the (partial) presence of the normal distribution, we use the non-parametric 2-sided Mann Whitney test for comparison. 2D kernel density estimation (KDE) is used to gain insight into potentially interesting patterns in 2D distributions in addition to the regression lines.

### Runtime and memory constraints and its consequences for parameters

The detection algorithm uses a window of size w, for 3D leading to a total window size of (2*w+1)^3^ voxels. For our results we set w to 2, so each window is 125 voxels large. Let N = X x Y x Z, the dimensions of both channels, and M = (2*w+1)^3^. Time complexity is dominated by the Spearman correlation, which needs to be applied N times, leading to O(N M log M) complexity. Space complexity is linear in N, however, to offer the end user maximal interpretability we output multiple intermediate stages, leading to an estimated memory use of ∼ 10 * N. For our data, depending on the cell size, memory usage averages between 32 and 128GB. The complexity analysis is essential to illustrate why we set w=2. A window size determines directly the minimum observable Spearman response. Larger windows therefore can detect fainter responses. However, a large window risks including multiple interaction patterns, for example, when an interaction zone is enclosed by 2 mitochondria. In such cases the response of both would cancel out the desired response, and we would lose information. The complexity analysis shows that increasing w leads to a cubic increase in runtime. All of these reasons lead us to use a window of 2, spanning ∼ 350 nm.

### Limitations

It is important to frame the proposed contribution within its limitations. First, any misaligned or unregistered channels can induce false responses or, more likely, cause contact detection to fail. Normalization of both input channels can destroy reconstruction and is not needed, as each stage is already adapting to the relative intensity distribution. Similarly, if the deconvolution is not calibrated correctly to match the empirical PSF, reconstruction can be compromised. Second, the recovery of sub-precision information is only possible because we exploit the local differential intensity profile of each voxel over a given window. If the signal to noise ratio is too low, differential analysis can break down. If the precision of the system decreases, or spans many voxels, the localization of the interface will become more challenging. For example, from Figure 2B it can be deduced that if the two intensity profiles widen too much, the negative correlation vanishes when their negative 2^nd^ differentials no longer overlap. This phenomenon will become more likely when object size decreases with respect to the width of the interface. For instance, while MCS-DETECT can in principle be applied to diffraction limited samples, recall of interfaces would be less precise. As shown in Figures 3, 4 and 5, two-fold improved resolution of 3D STED image stacks effectively enabled detection of MERCs, that present a 3-10 fold reduced width compared to the resolution of the system. Similarly, a too large window size can destroy a response, for instance when the window spans the size of both objects. In this case positive and negative differentials cancel each other out. Finally, anisotropic precision leads to responses that are less precise in the axis where precision is lowest. In our case precision is worst in Z, so recovered contact sites on top of mitochondria are likely to be captured with reduced precision. Finally, note that MCS-DETECT does not separate proximate contacts that a user could segment as two or more individual adjacent contacts. In future work a more refined approach will tackle the per-region identification even for proximate contacts, but this is limited by ground truth voxels annotated for specific contacts.

