## Supplemental Figures and Legends for "Membrane contact site detection (MCS-DETECT) reveals dual control of rough mitochondria-ER contacts"

### SUPPLEMENTAL FIGURE AND VIDEO LEGENDS

**Supplemental Figure 1. Parameter study of the proposed method.** A) We test 2 representative HT-1080 and COS-7 cells, with known distinct contact types. We vary the analysis window, a 3D cube of 5k x 5k x k over the mitochondria and contact channel. a) the surface coverage ratio stays stable for a range of k. b) The user can set a significance and statistical power threshold. This increase the minimum significance at which voxels can be detected, as well as the minimum correlation considered to be observable. As expected, when this threshold increases, expected differences between 2 representative cells decrease, as does the overall number of correlation voxels. At the limit of 100% confidence, no information would be left. To avoid false responses by bleed-through of signal in nearby Z-planes, as well as high background intensity in low SNR conditions, we apply an adaptive threshold in z-space. c) We observe that at low values, the inclusion of false responses masks any differences (z=1.5). At high values (3.5) the ER channel was visually degraded, we see that after z=3 the difference between the 2 cells is maximal and converged. B) The full reference algorithm pseudocode listing of each stage of the proposed method, enabling reproduction in any implementation. The actual Julia implementation used adds non-algorithm stages to deal with parallelization, optimization, error handling, and recording intermediate stages, which are out of scope for the purposes of this listing. Source code is being prepared for release under AGPL v3 license.

**Supplemental Figure 2. Identifying and filtering mitochondria.** A) For each contact, size and mean intensity of the adjacent mitochondria is plotted for two replicates, to indicate consistency across replicates. Note that this is by default larger than a segmentation method would compute. The size and mean intensity of mitochondria in HT-1080 and COS-7 cells present a clearly separable group of small, low intensity mitochondria structures. To report results on what are clearly and unambiguously mitochondria, corresponding to the mitochondria observed by EM, mitochondrial structures smaller than thresholds 9 (ln size) and 0.2 (mean intensity) were eliminated. B) 3D STED mages show mitochondrial labeling of HT-1080 and COS-7 cells before and after mitochondrial filtering (Bar = 10 μM).

**Supplemental Videos 1 & 2** Videos depict 360° views of 95$^{th}$ quantile MERCs in HT-1080 cells (Fig. 5C). MERCs display a high degree of complexity with multiple branch points and extending over several Z slices. Additional channels are rendered after each rotation in the following order: MERC channel (white), transparent mitochondria (pink), opaque mitochondria (red), and transparent ER (green).

**Supplemental Videos 3 & 4** Videos depict 360° views of 95$^{th}$ quantile MERCs in COS-7 cells overexpressing the OMM-ER linker (Fig. 5C). Planar MERCs follow the mitochondria in a single direction with limited travel to additional Z slices. Additional channels are rendered as described in the legend to Supplemental Videos 1 and 2.

**Supplemental Videos 5 & 6** Movies depict 360° views of $95^{th}$ quantile MERCs in COS-7 cells overexpressing WT Gp78 (Fig. 5C). MERCs display a similar phenotype to MERCs observed in HT-1080 cells. Additional channels are rendered as described in the legend to Supplemental Videos 1 and 2.

A

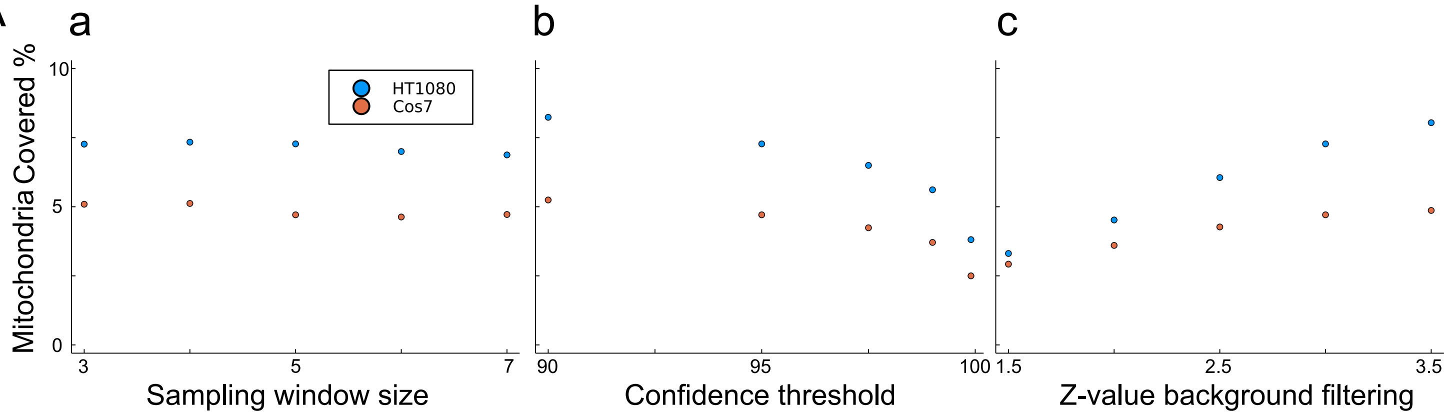


B

**Algorithm 1** Contact detection algorithm

**Input** $R, G$ : 2x3D, window size $w, \alpha, \beta, \sigma = 1$
**Output** $C_d$: contact prediction, confidence map $M_d$
$R_l, G_l \leftarrow$Gaussian(Laplacian($X$), $\sigma$) for X in $[R, G]$
$C_i, M_i \leftarrow$ spearman$(R, G)$ ▷ Alg.3
$C_d, M_d \leftarrow$spearman$(R_l, G_l)$
$C_d$ =sigfilter$(C_d, M_d, w, \alpha, \beta)$ ▷ Alg.4
$C_i$ =sigfilter$(C_i, M_i, w, \alpha, \beta)$
$C_d \leftarrow C_d.*(C_d. \wedge C_i)$
$C \leftarrow C_d.*$gradientfilter$(R_c, G_C)$ ▷ Alg.6
Return $C_d, M_d$

**Algorithm 2** Background removal

**Input** $R, G$, background filter strictness z
**Output** $R', G'$
$G'[x, y, z] \leftarrow (G'[x, y, z] > \mu_G + z * \sigma_G)$ ? 1 : 0
$R'[x, y, z] \leftarrow (R'[x, y, z] > \mu_R + z * \sigma_R)$ ? 1 : 0
Return $R', G'$

**Algorithm 3** Spearman

**Input** $R, G$, window size $w$
**Output** Correlation $C$, Confidence $M$
$N \leftarrow (2 * w + 1)^3$
**for** $x, y, z \in dim(R)$ **do**
$a, b \leftarrow R[x \pm w, y \pm w, z \pm w], G[x \pm w, y \pm w, z \pm w]$
$si_a$ =index(sort($a$))
$si_b$ =index(sort($b$))
$r = \frac{cov(si_a, si_b)}{\sigma_{si_a} \sigma_{si_b}}$
$C[x, y, z] = |\text{Min}(r, 0)|$
$M[x, y, z] = \sqrt{\frac{N-3}{1.06}} *$atanh$(r)$ ▷ z-score of Fisher transform
**end for**
**Return** $C, M$

**Algorithm 4** Significance Filter

**Input** $X$, window size $w, \alpha, \beta, M$
**Output** Filtered $X'$
$r_\beta$ =minr$((2 * w + 1)^3, \alpha, \beta)$ ▷ Alg.5
$X'[x, y, z] = X[x, y, z] * (M[x, y, z] < \alpha \wedge X[x, y, z] > r_\beta) \quad \forall x, y, z \in dim(X)$
**Return** $X'$

**Algorithm 5** Minr

Input $N$ (sample size), $\alpha, \beta$
Output Minimum observable correlation r
$N' = \infty$
$z_\alpha, z_\beta = qnorm(\alpha, \beta)$▷ Quantile of normal distribution
**for** $r \in [0, 1]$ **do**
z' $= \frac{1}{2} \log \frac{1+r}{1-r}$
N' $= \left(\frac{z_\alpha + z_\beta}{z'}\right)^2 + 3$
**if** $N' < N$ **then**
**Return** r
**end if**
**end for**
**Return** 1

**Algorithm 6** Gradient filter

**Input** $R, G$
**Output** Mask where gradient detects no artifacts
$R_d, G_d \leftarrow |\nabla_R^3|, |\nabla_G^3|$ ▷ 3rd derivative intensity
**Return** $\neg((R_d. * G_d. = 0). \wedge (R_d. + G_d.! = 0))$

**Algorithm 7** Analysis

**Input** $C, X, Y, Z$, minimal observable size $P$
**Output** Contacts, Features
$C' \leftarrow$ morphological closing(C)
$O \leftarrow$ connectedcomponents(C)
$O' \leftarrow \{o \mid |o| > P \quad \forall o \in O\}$
**for** $i \in dim(C)/(X, Y, Z)$ **do**
$cube \leftarrow O[x + iX, y \pm iY, z \pm iZ]$
result[i] = analyze(cube)
**end for**
**Return** result, $O$

Supplemental Figure 1

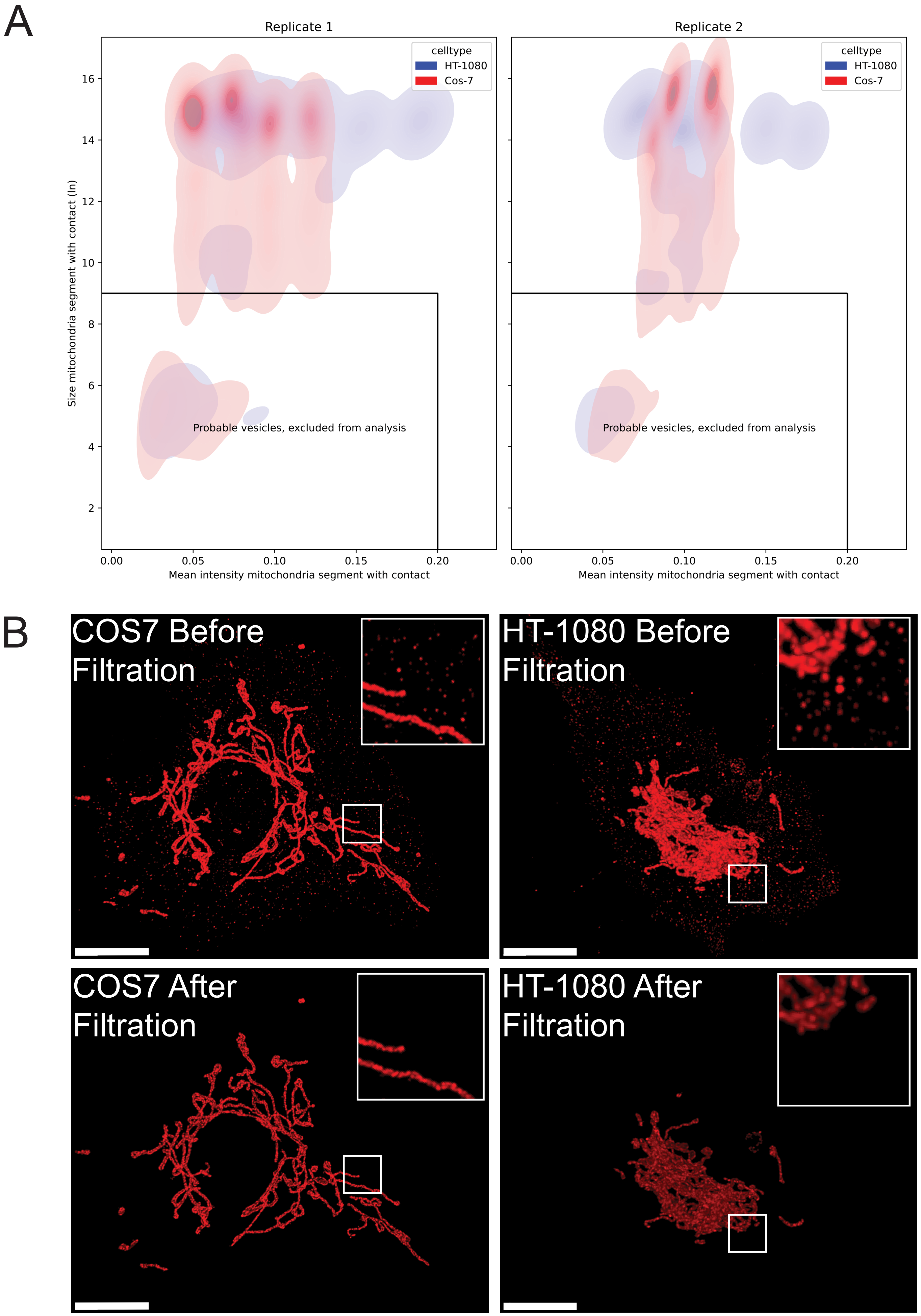


Supplemental Figure 2
